# Local decondensation at double-stranded DNA breaks modifies chromatin at long distances and reduces encounter times during homology search

**DOI:** 10.1101/508689

**Authors:** A. Amitai, O. Shukron, A. Seeber, D. Holcman

## Abstract

Double-strand break (DSB) repair by homologous recombination (HR) requires an efficient and timely search for a homologous template. Here we first study global chromatin re-organization following a single DSB: due to the potential release of cross-linkers such as cohesin and CTCF molecules near the DSB site, loops are released and chromatin is decondensed, explaining the change of chromatin locus motion at larger genomic distances. This mechanism provides an elementary explanation for the increase of the anomalous exponent at sites located far away from the DSB, after break induction. Second, we explore the consequences of chromatin reorganization for the homology search during DNA repair: using polymer models, we estimate the mean first encounter time (MFET) between two loci on the chromatin in a confined nucleus. Reducing tethering forces, as reported experimentally on chromatin, is associated with a local de-condensation near the break followed by the extrusion of the breaks. Consequently, we report here that the mean first encounter time between homologous sites is decreased by two orders of magnitude even when the homologue sequence is located on the nuclear boundary. To conclude, our results suggest that local changes in inter-nucleosomal contacts near DSBs, by cohesin removal, remodel the chromatin and drastically shorten the time required to complete a long-range search for a homologous template.

## 1 Introduction

DNA double-strand breaks (DSBs) can be repaired by one of two major mechanisms: non-homologous end joining (NHEJ), whereby the broken ends of the DNA are simply religated [1], or homologous recombination (HR), which, if the sister chromatid is likewise damaged or unavailable, entails a physical search by the broken strands for a homologous DNA template. This template is then used to repair the break [2, 3]. The homology search (HS) occurs in budding yeast even when the template for repair is located on another chromosome, yet HS appears to be the rate-limiting step [4–7] a general mechanism of repair across species but the physical mechanism remains unclear. Given that base-pairing occurs on an atomic level where DNA is folded into chromatin fibers of 10-300 nm [3] in a nuclear volume that contains millions of potential pairing partners, the search for homology must occur on multiple spatial-temporal scales, extending from the molecular to the nuclear level. However, it is not clear how local changes in protein-DNA or DNA-DNA interactions affect long-range movement, nor is it clear how local and long-range dynamics are linked. These dynamics must be resolved [6–10], if we are to understand the physical basis of this phenomenon.

Chromatin is constantly moving inside the nucleus, driven by collision, in a sub-diffusive motion, that can be characterized by the dynamics of a genomic locus [11]. These stochastic dynamics, modulated by active ATP-dependent mechanisms [12, 13], are physically described by four independent parameters: the effective diffusion coefficient *D*_*c*_, the anomalous exponent *α*, the overall tethering force spring constant *k*_*c*_, and the length of loci confinement *L*_*c*_ [10, 11]. Chromatin movement could generate random encounters between different genomic loci, such that two distant sites for sequence-specific binding factors are able to interact, generating DNA loops that bring enhancer and promoter together in a transient but efficient interaction [14,15]. The rate of genomic looping depends on key parameters such as size of confinement [16], chromatin folding [17], and can be impeded by sites of tethering [11].

Recently we reported [11, 18] that a change in the dynamics and organization of chromatin following DSB induction affects the rate of interaction between two non-contiguous sites. By developing polymer models [19–21], that account for the statistics of single particle trajectories, we found that a DSB is associated with a local chromatin decompaction, characterized by an increase of the anomalous exponent *α*, a release of local tethering forces, and an expulsion of the broken site to the periphery of the chromatin globule. How these changes affect long-range motion of chromatin at a DSB remain unexplored.

Using polymer simulations, we interrogate here the consequences of chromatin re-organization following DSB on the search time for a template. In particular, we explore using coarse-grained polymer models, how a single chromosomal locus can explore the entire territory of another chromosome, or the entire genome, and to find a small target sequence for strand invasion. Despite the need to scan millions of base-pairs for the correct sequence, the search for a homology in yeast can occur within a few hours [4, 22]. This rate is all the more remarkable if one considers that the process is driven by diffusion, and must overcome physical barriers and repulsion forces during the invasion of a chromatin domain. We find here that the extrusion of the affected monomers reduces the mean first encounter time of the two broken ends by two orders of magnitude, even when the homologue sequence is located on the nucleus boundary. Finally, we predict that local expansion of the chromatin at the site of a DSB is a key step of HR. The present results provide a scenario and prediction that should be tested about chromatin reorganization during HR in the nuclear environment.

## 2 Results

### 2.1 Chromatin remodeling near a break can modify chromatin dynamics far from a DSB

The induction of a single DSB can lead to a global chromatin reorganization [23, 24]. Such re-organization was probed by SPTs of an undamaged single tagged locus, located far from a DSB. Surprisingly, loci motion is affected by induced breaks, characterized by an increased anomalous exponent [18] suggesting a local chromatin decompaction post DSB. The reasons for such long-distance effect remains unclear, because it is not clear how local decondensation of chromatin can influence the dynamics of sub-nuclear regions located far away from the DSB. Since DSB induction is associated with local nucleosome repositioning [25] and removal of cross-linkers such as cohesin or CTCF, to explore the possibility that removal of these cross-linker alone can lead to long-range changes, we used here a randomly cross-linked (RLC) polymer model (Methods 5.1). Removal of cohesin in G1 cells (no sister chromatid present) resulted in increased loci motion. This implies that cohesin could be a cross-linker in yeast [26]. Our goal is to test the hypothesis that few cross-linkers molecules generate long-range loops [27]. Thus, By removing these cross-links at the vicinity of a DSB, loops are destabilized and, consequently, results in a local decompaction around the DSB by cross-linker removal. We shall see that this removal has long-range consequences.

Using the RCL polymer model [20] (see subsection 5.1), we examine three scenarios of removal of cross-links following a single DSB. First, *N*_*r*_ ∈ [0, 45] cross-links are removed uniformly at random following DSB induction (Fig. 1A left); Second, only the *N*_*r*_ ∈ [0, 45] closest cross-links to the DSB are removed, starting with those near the break and then further away in an increasing manner (Fig. 1A center). Third, we remove all cross-links located within a distance of 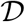 ∈ [0.03, 0.2]*μm* from the DSB, (Fig. 1A right). We performed 1000 simulations for each *N*_*r*_ ∈ [0, 45] and 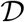 ∈ [0.005, 0.2]*μm* of total time of 33 seconds post DSB (parameters described in Table 1). We then generated trajectories for all monomers to characterize their motion by computing four statistical parameters [28]: length of confinement *Lc* (Eq. 8), effective diffusion coefficient *D* (Eq. 9), spring constant *k*_*c*_ (Eq. 10) of the effective tethering force, and the anomalous exponents *α* (Eq. 7), as the number of removed cross-links *N*_*r*_ or the distance 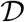 increases.

**Table 1:** Parameters used for simulation of a DSB in the RCL polymer

| parameter | value | description |
| --- | --- | --- |
| $N$ | 100 | Number of monomers |
| $b$ | $0.2 \mu m$ | STD of connectors' length |
| $D$ | $0.008 \mu m^2/s$ | Diffusion coefficient |
| $\Delta t$ | $0.0033$ s | Simulation time step |
| $N_s$ | 10000 | Number of steps after relaxation |
| $N_c$ | 50 | Number of added connectors |
| $N_r$ | $[0, 45]$ | Number of connectors removed following DSB |
| $\mathcal{D}$ | $[0.0050.2]\mu m$ | Maximal distance to disconnect cross-links |

**Figure 1:**
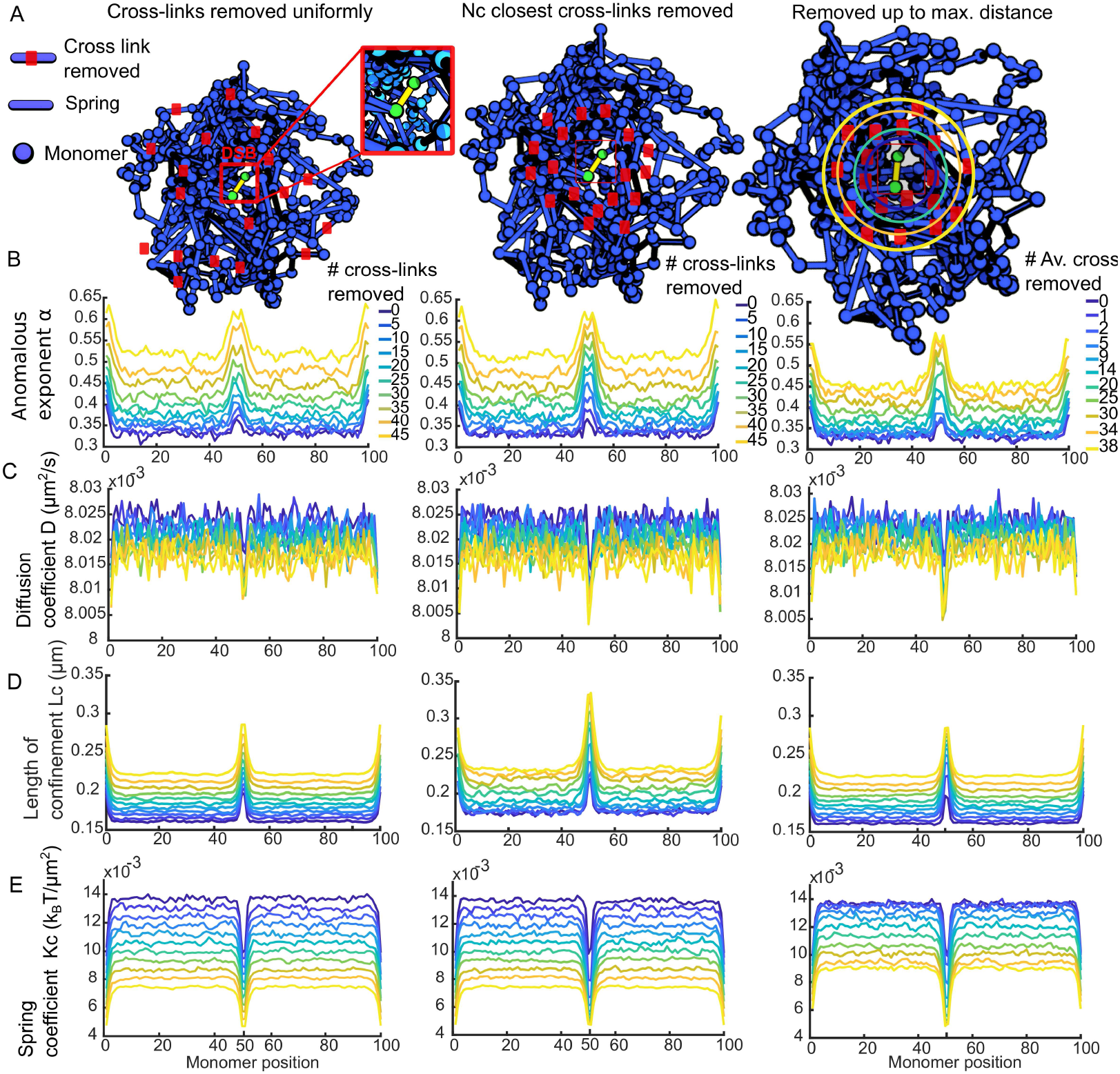
Chromatin reorganization following a single DSB. **(A)** Following a DSB between central monomers 50 and 51 (yellow, inset), three scenarios are simulated using a RCL polymer model (blue), where *N*_*r*_ cross-links are removed: uniformly (left), then the *N*_*r*_ ∈ [0, 45] closest cross-links (center), and then up to maximal distance 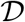 ∈ [0.005, 0.2]*μm* (left). Removed cross-links are in red. **(B)** Anomalous exponent *α* (Eq. 7) after DSB induction for uniform (left), distance-based (center) and max distance (right). **(C)** Effective diffusion coefficient *D*_*c*_ (Eq. 9) for the three cases as in panel B. **(D)** Length of confinement *Lc* (Eq. 8). **E.** Spring coefficient of the tethering forces *Kc* (Eq. 10).

We found that near the break induced between monomer 50 and 51, the anomalous exponent increases from 0.35 when no cross-links are removed, as shown in Fig.1B (blue) to 0.6 in Fig 1B (yellow) when 45 cross-links are removed for all three scenarios. The anomalous exponent of monomers far from the break increases from 0.3 to 0.5 when 0 and 45 cross-links where removed post DSB, respectively. This increase in *α* is associated with a local decompaction of the chromatin around a DSB. The effective diffusion coefficient remained fixed at 8.010^−3^*μm*^2^/*s* for monomers around the DSB, for all three scenarios (Fig. 1C. In addition, the diffusion coefficient slightly increased from 8.010^−3^ to 8.10^−3^*μm*^2^/*s* for monomers located far away from the DSB, when 0 to 45 monomers where removed post DSB, for all three scenarios, respectively. The length of constraint *Lc*, before and after a break, increased from 0.21*μm* to 0.27*μm*, near the break, for the cases of uniform and maximum distance based removal of cross-links, whereas for the distance-based removal, *Lc* near the break increased from 0.23 to 0.33*μm*, when *N*_*r*_ = 0 to 45 cross-links were removed, respectively. For monomers far from the break, we observe a change from 0.13 to 0.16*μm* when 0 to 45 cross-links were removed post DSB, respectively (Fig. 1D). The effective spring coefficient *Kc* showed similar behavior for all three scenarios, reducing from 14.10^−3^ to 7.10^−3^*k*_*B*_*T/μm*^2^ when 0 to 45 cross-links were removed following DSB.

We then examined the average (over monomers) of the statistical parameters computed post DSB for all three scenarios of removal of cross-links (Fig. 2). We found that the average anomalous exponent 〈*α*〉 increased from 0.34 to 0.53 fr the cases of uniform (blue diamonds) and distance-based (red squares) removal of cross links (Fig. 2A), when 0 to 45 cross-links were removed post DSB, respectively. For the case of maximal distance-based removal of cross-links the average anomalous exponent (yellow circles) increased from 0.34 when no cross-links were removed, to 0.46 when 38 cross-links were removed on average. The average effective diffusion coefficient 〈*D*〉 remained constant at 8.010^−3^*μm*^2^/*s* when 0 to 50 cross-links were removed for all three scenarios. The average length of confinement 〈*Lc*〉, showed similar behavior for all three scenarios (Fig. 2C), increasing from 0.16 to 0.23 *μm* when 0 to 45 cross-links were removed post DSB and when 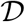 increased from 0.005 to 0.2 *μm*. Finally, the average effective spring coefficient 〈*Kc*〉 decreased similarly in all three scenarios from 1.3510^−2^ to 0.710^−2^*k*_*B*_*T/μm*^2^ when 0 to 45 cross-links were removed following DSB.

**Figure 2:**
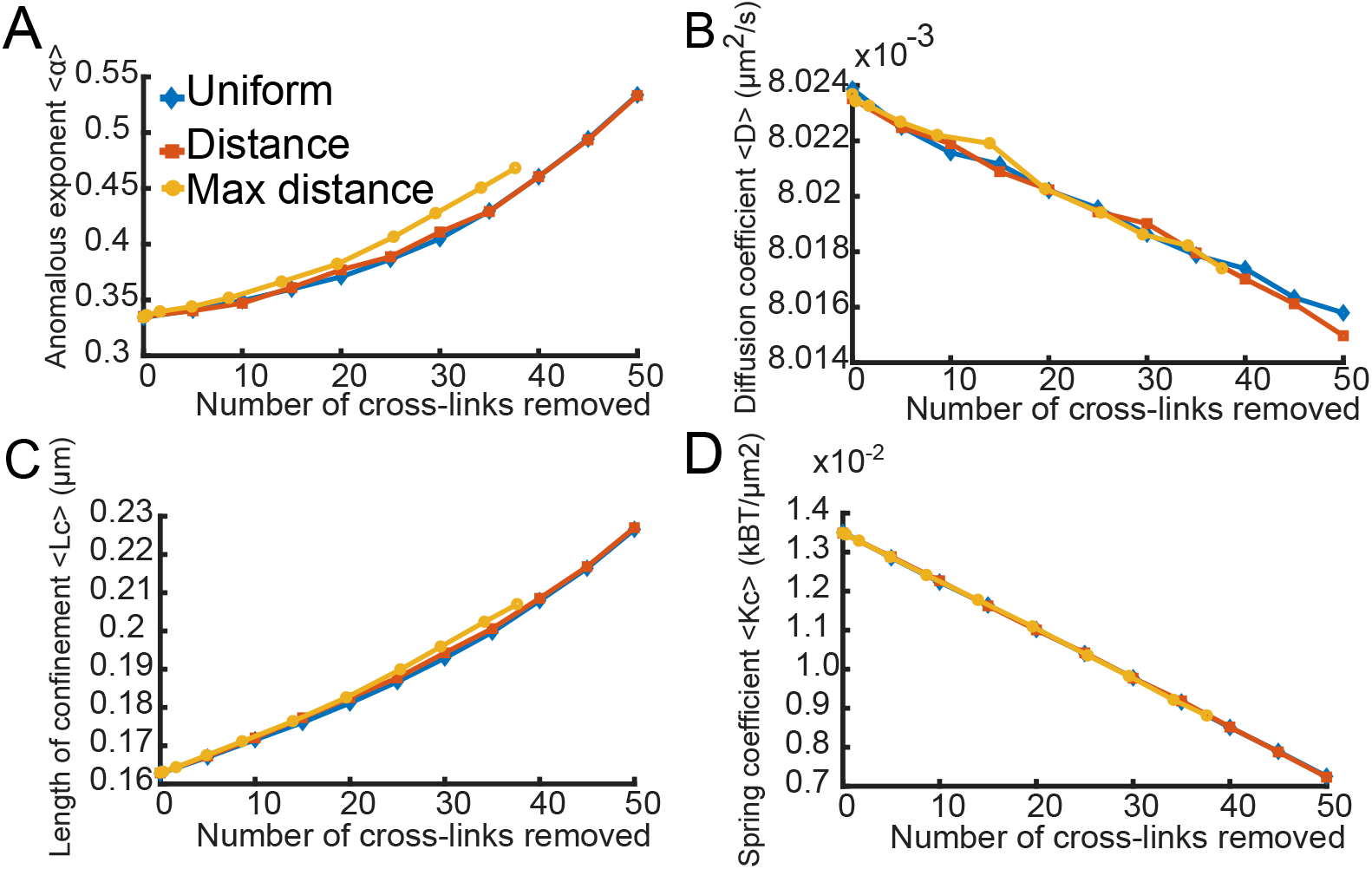
Averaged parameters vs number of removed cross-linked for chromatin reorganization following DSB. **A.** Average anomalous exponent 〈*α*〉 computed over all monomers from Fig. 1B, for the three scenario of removing cross-links: uniform (blue diamonds), distance-based (red squares), and removal up to maximal distance (yellow circles). **B.** Average apparent diffusion coefficient 〈*D*_*c*_〉 computed from Fig. 1C. Color scheme as in panel A. **C.** Average length of confinement *Lc* computed from Fig. 1D. **D.** Average apparent spring coefficient *Kc*, computed from Fig. 1E.

To conclude, we found that local removal of cross-links around DSB led to a large-scale decompaction of chromatin, characterized by an increase of the anomalous exponent *α* and the length of constraint *Lc*. Local removal of cross-links changes the statistics of the four variables (Fig. 2) and in particular cross-link removal can generate long-range loops (see also [29]). In this sense, few cross-linkers positioned at a strategic location that attach various chromatin fibers control chromatin dynamics even far away.

## 3 Time scale of homologous search for a relocalized DSB

If chromatin expands around a DSB, with long-range consequences, these changes in chromatin at the site of a break can also affect the medium-range architecture of chromatin near the break. Using numerical simulations of the *β*-polymer model [19], we explored the local structure the chromatin by reducing the intrinsic interactions of a single monomer, representing a local release at the site of the DSB. A *β*-polymer (for 1 < *β* < 2) is characterized by long-range forces between monomers responsible for the polymer condensation level. While *β* = 2 is the classical Rouse chain, a smaller value of *β* leads to highly compacted polymer. This polymer model accounts for the forces among monomers, when the anomalous exponents *α* is prescribed within the range [0 − 0.5]. We interpret values of *α* larger than 0.5 as the consequence of an additional external deterministic forces acting on the chromatin and the intra-chromatin forces. To account for changes in chromatin structure and to avoid interpenetration of the polymer model representing chromatin, we added repulsion interactions between each monomer i.e., Lennard-Jones (LJ) interactions to the polymer model. We use a polymer of length *N* = 33 monomers, with the coefficient *β* = 1.5, which results in a highly connected polymer (Fig. 3). With the addition of LJ-interaction forces, the relationship of the beta-polymer no longer holds [21].

**Figure 3:**
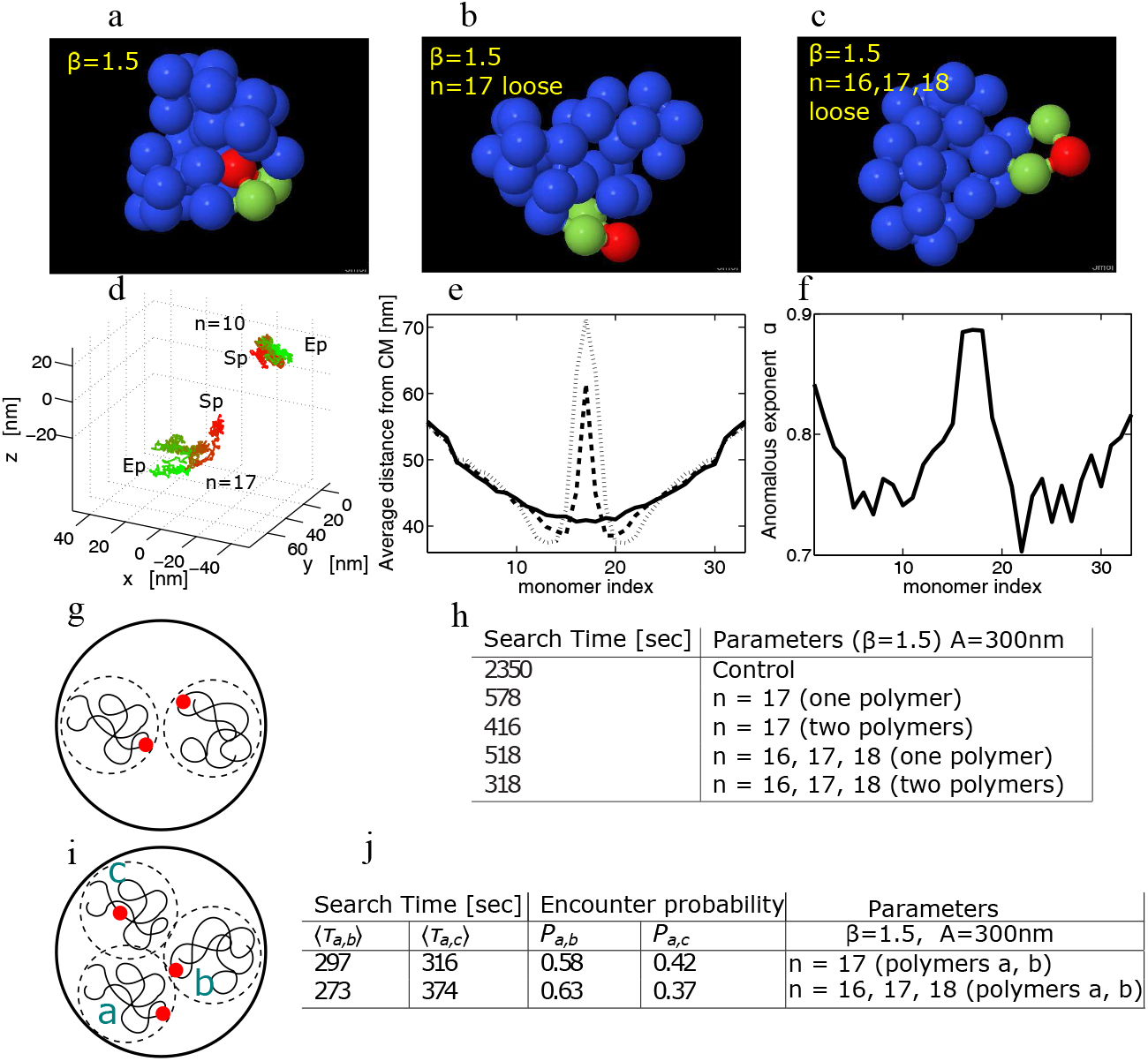
Extrusion of a broken locus at the periphery of a chromatin domain. (**a**) Steady-state configuration of a polymer model (*β* = 1.5 and *N* = 33 with Lennard-Jones interactions), showing the middle monomer *n* = 17 (red) and its two neighbors *n* = 16, 18 (green): in the control condition (a), (**b**) long-range interactions of the *β*−polymer are removed at location *n* = 17 (**c**) and at the three monomers *n* = 16, 17, 18 (c). (**d**) Two trajectories of monomer *n* = 10 (no changes in local interactions) and for *n* = 17 (interactions are removed) with respect to the CM of the polymer. The initial position is marked by the start point *S*_*p*_ (red), the long-range interaction for monomers *n* = 16, 17, 18 are instantaneously removed. From 0s to 0.05s, the color changes gradually to green until the end point *E*_*p*_. (**e**) Average distance of each monomer from the CM corresponding to (a) (full line), (b) (dashed line), (c) (dotted line). (**f**) Average anomalous exponent *α* for different monomers during the extrusion process for case (c). (**g**) Search process between two extruded monomers located at the periphery of their chromatin territories. (**h**) MFET of two middle monomers *n* = 17 of each polymer represented in (g) when the interaction between monomers are not perturbed (control a), the polymer interactions on a single polymer acting on the middle monomer are disrupted and then on both of them (b) and finally, when the forces are disrupted (case c) for one and then both polymers. Polymers are initially placed in a sphere of radius *A* = 300*nm* and the encounter radius is 5*nm*. (**i**) Competitive search between extruded monomer from polymer *a* and the middle monomers of either polymer *b* or *c*. The middle monomer of *b* is extruded. (**j**) MFET and the encounter probabilities of *a* with *b* or *c* for the competitive searches.

To test the effect of the DSB on chromatin organization, we reduced the interactions of the middle monomer (red monomer Fig. 3), to only include nearest-neighbors interactions (green monomers, Fig. 3b). Doing so we accounted for the eviction of nucleosomes adjacent to a DSB. Surprisingly, this local modification had a dramatic effect on the global structure of the polymer and the spatial distribution of the middle monomer: removal of the long-range forces of the middle DSB led to its extrusion to the periphery of the polymer’s globular shape (Fig. 3b). Chromatin decompaction was obtained after the removal of local forces acting on the two neighbors of the middle monomer 17-18 (Fig. 3c). The monomer extrusion is persistent and irreversible, so that the extruded monomer remains at the periphery of the chromatin domain, suggesting that a following DSB the damaged chromatin is extruded from the sub-chromatin region where it is embedded.

We followed over time the extrusion event and measured the displacement of both unmodified monomers and the DSB modified monomer (Fig. 3d). While the unmodified monomer did not change its position significantly during polymer reorganization, the modified monomer exhibits directed motion from the center to the periphery of the domain (Fig. 3). To quantify this effect, we plotted the average distance of each monomer from the center of mass (CM) of the polymer: Fig. 3e shows that before the removal of forces, the middle monomer was the closest on average to the CM, while after local relaxation, it became the most distant. To check whether or not the anomalous exponent *α* was modified during the extrusion event, we computed it for each monomer during this process (Fig. 3f). For the middle monomer *α* changes from 0.52 to 0.88, suggesting that during extrusion, DSB motion is driven by a deterministic force generated by remaining monomers, corresponding to the undamaged chromatin region.

### 3.1 Reduced local monomer interactions and extruded DSB monomer decreases Homology Search time

To examine the consequences of reducing the intrinsic local interactions and the extrusion of the DSB middle monomer on Homologous Search, we estimated the time for two monomers located on two separated *β*-polymers (with *β* = 1.5) to contact for the first time (Fig. 3g-h). When the two monomers are not extruded (control), the MFET computed from simulations is 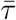 = 2350*s*. If one monomer was extruded, then the mean encounter time reduces to 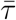 = 578*s*. We also accounted for the observations that a break and its template show increased movement [8] following DSB. Multiple DSBs cause a general increase in chromatin mobility [23]. Thus we also extruded the homologous template, and in that case, the MFET reduces to 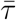 = 416*s*.

To confirm the observed reduction of the search time, we released the forces of the neighboring monomers 17 and 18 which led to a similar reduction in MFET (Figure 3h). When we reduced the intra-connecting forces on both polymers, we observe further drop in the MFET, mediated by the collective self-avoiding interactions between monomers that generate an effective energy barrier around the middle monomer. This barrier hampers contact with other monomers or polymers, acting as a physical barrier. Thus, similar to the energy barrier that enzymes have to overcome for activation, we propose that an effective screening potential is the energy required for a chromatin locus to explore another chromosome territory. This value may depend on the position of the monomers within their territories, the forces between monomers and on polymer length. This effect was previously anticipated in [30] for a polymer modeled as a soft colloid.

In summary, a decrease in the search time (MFET) reflects a reduction in the strength of the effective screening potential that the searching monomer experiences. This suggests that encounters between two monomers require activation over a potential barrier [31]. Since the middle monomer is the closest one to the center of mass, it is the most shielded from external contacts. Upon extrusion to the periphery of the local chromatin domain, the strength of the potential decreases, facilitating HS and leading to a 4.5-fold reduction in search time for a monomer adjacent to the extruded DSB. If both polymers are reorganized, the monomers of the two will come into contact 7.5-fold faster (Fig. 3h).

### 3.2 First encounter time when a DSB competes for two different repair sites

When a DSB recombines with an incorrect sequence it generates a translocation, which in the worst of cases misregulates genes and contributes to oncogenic transformation [32]. We therefore explored a scenario in which we model a potential translocation event between two polymers with a third polymer present. We tested the effect of DSB extrusion on the MFET and encounter probability between the middle monomers of three polymers (*a, b, c*) where *a* contains the DSB, and *b* and *c* represents template sequences. An encounter with *c* represents a translocation, while an encounter with *b* represents a correct search outcome (Fig. 3i).

The middle monomers of polymers *b* and *c* are competitors for the interaction with the broken site *a*. If the middle monomers on polymer *a* and *b* are extruded, the MFET is *τ*_*ab*_ = 297*s*, while *τ*_*ac*_ = 316*s* and the respective encounter probabilities are *P*_*ab*_ = 0.58 and *P*_*ac*_ = 0.42. When the internal forces are removed between monomers 16-17-18 in polymers *a, b*, as described in Fig. 4c, the difference is increased: *τ*_*ab*_ = 293*s* and *τ*_*ac*_ = 374*s* and the respective encounter probabilities are *P*_*ab*_ = 0.63 and *P*_*ac*_ = 0.37 (Fig. 3j).

**Figure 4:**
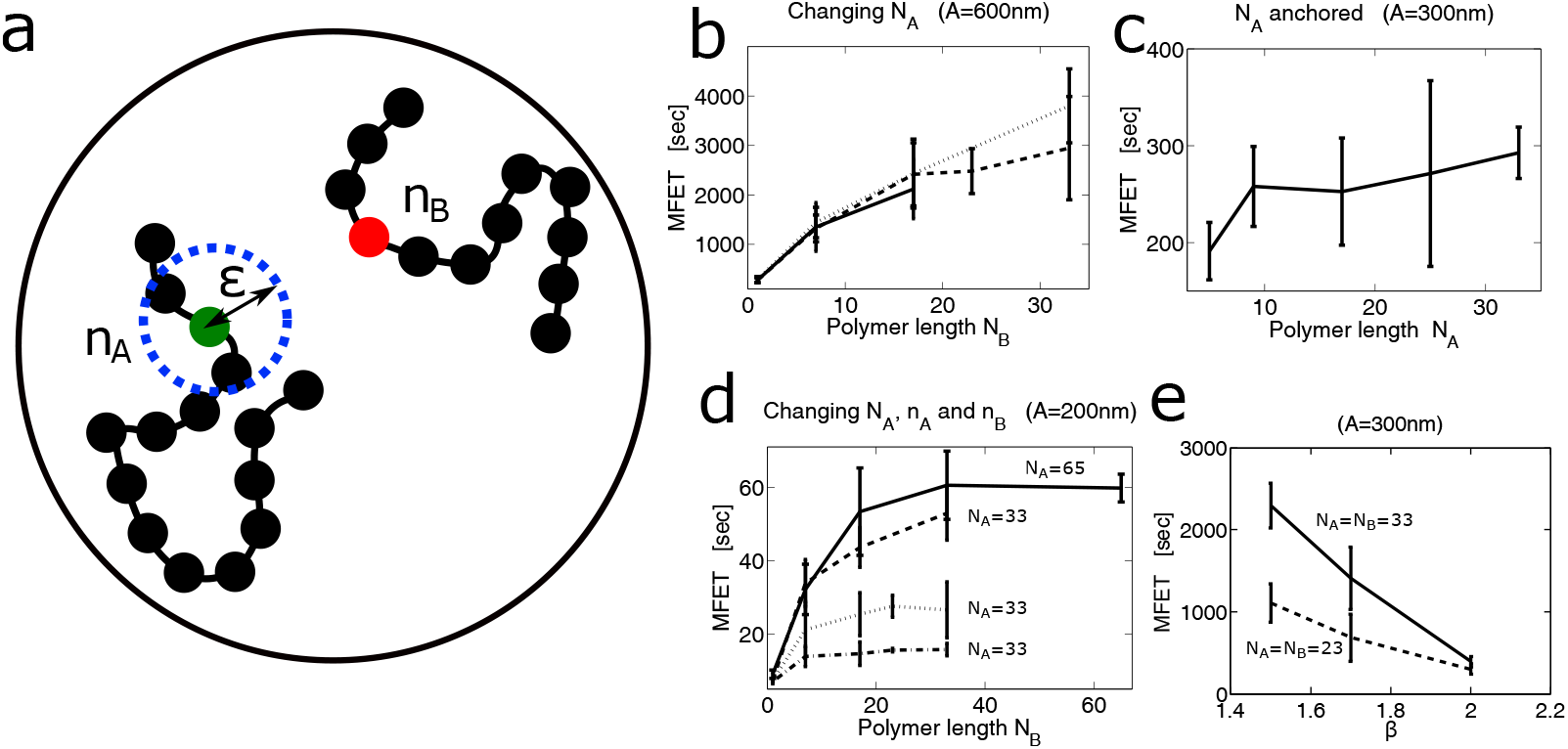
Search process for two monomers located on two self-avoiding polymers. (**a**) Illustration of two polymers in a nucleus. Monomers *n*_*A*_ and *n*_*B*_ interact when they are located within the activation radius *ε*. (**b**) MFET versus length polymer for two self-avoiding (SA) polymers in a sphere of radius *A* = 300*nm*. The length *N*_*B*_ of one polymer is the x-axis, while the different curves correspond to different values of the length of polymer *N*_*A*_ : 17 (full line), 33(dashed line), 65(point-line). Interacting monomers are the middle ones. (**c**) MFET versus polymer length *N*_*A*_ located in a sphere *A* = 300*nm*. Polymer *B* is a freely diffusing (with length *N*_*B*_ = 9), while the middle monomer of polymer *A* is anchored at the center of the sphere. (**d**) MFET versus polymer length for two self-avoiding polymers in a ball of radus *A* = 200*nm*. In the x-axis, we represented the length of one polymer B. The curves correspond to different values of polymer A for various length *N*_*A*_ and different monomers *n*_*A*_ and *n*_*B*_: interacting monomers are located either at middle of the chain (*N*_*A*_ = 65 solid line), (*N*_*A*_ = 33 dashed line), (*N*_*A*_ = 33 with interacting monomer on *A* and in the middle and on *B* at the end (dotted line), (*N*_*A*_ = 33 and both interacting monomers are at the ends, dotted line). (**e**) Encounter time of two middle monomers part of *β*-polymers with SA interaction of length *N* = 33 (solid line) and *N* = 23 (dashed line) in a ball of radius *A* = 300*nm*.

Therefore, a DSB that is extruded will more likely find a template that is also extruded. While it is unlikely that a cell can specifically increase the movement of the correct template sequence, it may favor sequences in which chromatin organization is altered by transcription or other domain modifying events, or it can alter the mobility of all chromatin in the nucleus. We suggest that a stochastic extrusion of undamaged loci throughout the genome in response to DNA damage may itself favor HS and facilitate repair [23, 33]. However, when there are several breaks, extruded domains may interact leading to an increase in translocation probability.

### 3.3 Search process for two monomers located on two self-avoiding polymers

In order to simulate how nuclear geometry, position of the DSB along the DNA strand and condensation of chromatin affect HS, we studied the MFET for two monomers located on two self-avoiding polymers (Fig. 4). We found that increasing the length of the polymer increased the MFET (Fig. 4b,c), while either reducing the condensation of the chromatin or positioning the DSB at the end of the polymer fig. 4d-e reduced MFET. We conclude that homologous search is influenced by polymer length, compaction and linear position of the break along the DNA fiber.

### 3.4 How relocation of DSB at the nucleus periphery affects HS

In budding yeast, persistent DSBs that do not have a homologous donor shift to the nuclear envelope [34–36], where they are anchored either at nuclear pores or at the SUN domain protein, Mps3. However, at the same time the DNA damage response causes a general increase in movement of undamaged loci [8, 23]. Therefore, we incorporated perinuclear tethering of one polymer into our modeling. To characterize the search process under these conditions, we scored the MFET between a DSB located on the boundary of a sphere (a single monomer not a polymer) Fig. 5a) and a *β*-polymer with self-avoiding interactions. We ran Brownian simulations of the polymers condensed to various degrees. We found that as *β* decreased, and the polymers became more condensed, the MFET for the peripheral target increased (Fig. 5b). This effect results from an effective screening potential barrier generated by the tethering of the monomer at the nuclear surface. In that case, the searching monomer needs to overcome this barrier as it approaches the nuclear surface.

**Figure 5:**
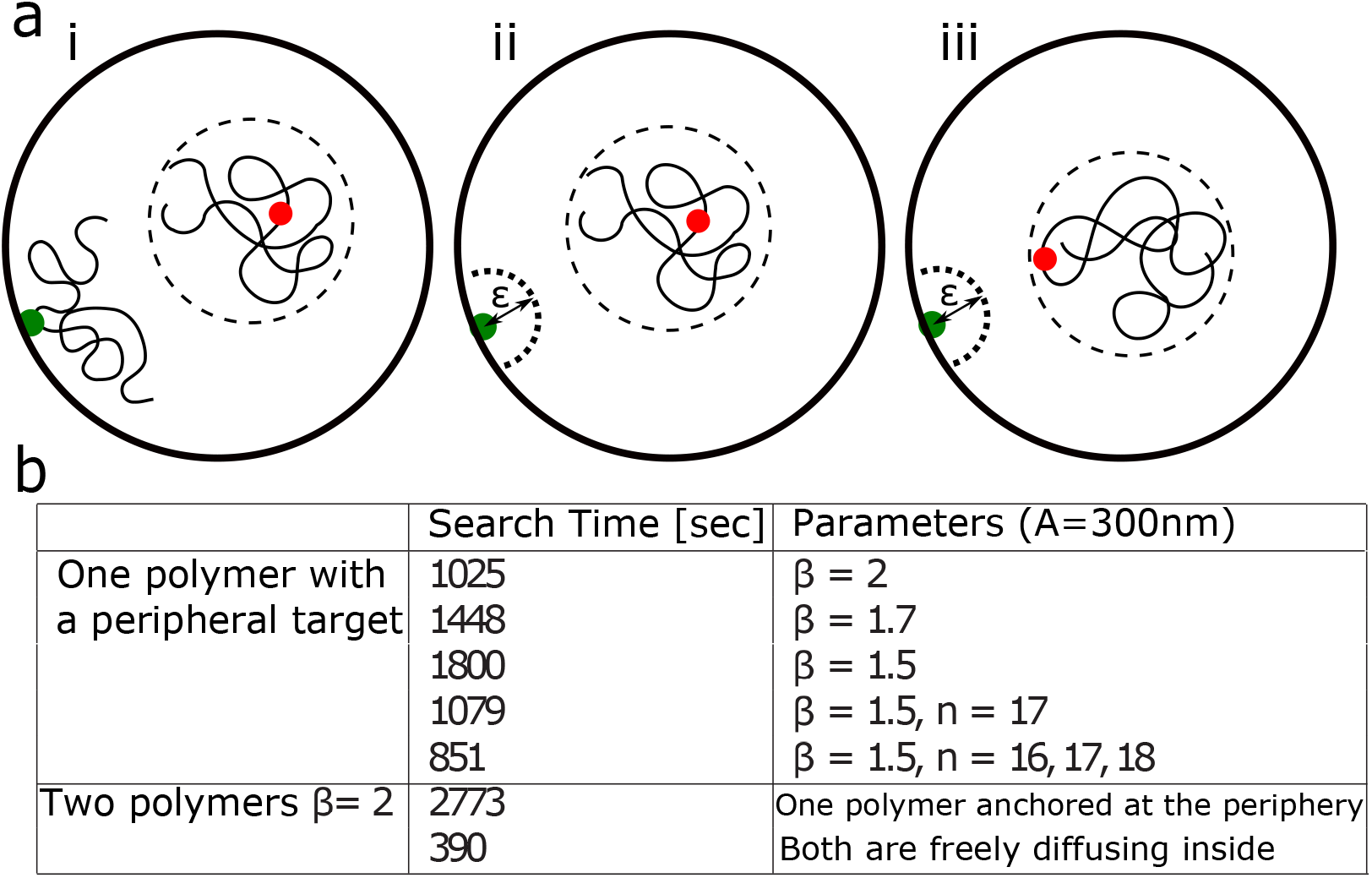
Polymer search process for a target monomer attached to the periphery. (**a**) (i): schematic representation of a polymer searching for a target (green) located on the nuclear envelope. The interacting monomer (red) is the middle one. (ii) Representation of a monomer (red) on one polymer searching for a monomer (green) part of another polymer attached at the nuclear envelope. The search ends when the two monomers are inside a ball of radius *ε* = 5*nm*. (iii) scheme of the polymer where the interacting monomer is extruded to the periphery of the polymer globule. (**b**) Summary of mean search time of the target in a ball of radius *A* = 300*nm* for different *β*−polymer of length *N* = 33 with LJ interactions. For *β* = 1.5, the long-range interactions are removed in two cases: at monomer *n* = 17 and *n* = 16, 17, 18. Finally, we show the cases where the target is the middle monomer and belong to another polymer (total length *N* = 33) attached at a distance 30*nm* from the nuclear envelope.

Finally, we simulated the search process when the DSB is the middle monomer of another polymer (*N* = 33), attached to the nuclear periphery, at a distance of few nm (Fig. 5f, right). As the DSB is now shielded by the other monomers of the chain, the MFET is larger than that of a break that was not a part of a polymer (2773s vs. 1025s), and the MFET for two freely diffusing polymers is even lower (390s vs. 2773s, Fig. 5g). This confinement effect suggests that searching for a target located at the nuclear periphery requires an extended search time. However, by loosening three local monomer interactions around the break, the search time can be reduced by a factor 2.5 (851s vs 1800s, Fig. 5g). We conclude that in order to facilitate the search by a DSB at the nuclear surface, both peripheral and target chromatin should be decondensed. This phenomenon reflects the additional barrier (screening potential) generated by the interaction of the monomer with nuclear envelope, leading ultimately to a search time larger than that for two free polymer chains.

## 4 Discussion

Recent progress in modeling of transient dynamics of polymers has illustrated how physical constraints define monomer motion, while allowing effective simulation of both intrinsic and extrinsic physical parameters. We applied these notions to chromatin organization in the nucleus. Several novel insights arise from this analysis. First, we found that looping induced by cross-linkers such as CTCF, cohesin or condensin create long-range connections. So that removing local cross-linkers can create large-scale reorganization, revealed by a change of loci dynamics (measured by the anomalous exponent). This result can be summarized as follows: long-range chromatin dynamics is controlled by the amount of loops and local cross-linkers (see also [**?**, 29]). Second, we confirmed that the motion of a single locus is characterized by an anomalous exponent, which depends in part on intrinsic polymer properties [19]: After the anomalous exponent has been extracted from single empirical trajectories, it is possible to construct a corresponding polymer that reproduces aspects of the measured dynamics both by contraining the number and position of cross-linkers. Analysis of such a polymer reveals how the sampling rate for imaging acquisition can affect the interpretation of the polymer motion [11]. Indeed, as the sampling time scale increases, more of the polymer length contributes to the motion, as do potential extrinsic forces that act over a longer time scales. Finally, we have explored here the consequences of chromatin reorganization on HS, investigating, using polymer simulations, the consequence of expelling the DSB to the periphery of sub-chromatin region, reposition the DSB at the nuclear periphery by removal of local forces. At this stage, we propose that a small region, that we call focal point, characterized by a high concentration of cross-linkers, controls long-range chromatin folding and dynamics. Removing cross-linkers around one of the focal point leads to chromatin decondensation and increase of local and long-range chromatin locus motion, leading to facilitate search during HS. It would be interesting to investigate the distribution of these focal points and to observe experimentally the consequences of removing them one by one on nuclear dynamics.

Previous studies of chromatin locus dynamics have been restricted to MSD analyses, which monitor the volume of sampled nuclear space, without shedding light on the nature of the forces acting on the locus in question. Such studies were able to document an enhanced sampling of the nuclear volume following DSB induction in response to the DNA damage checkpoint, but nothing more [8, 23, 37, 38]. Importantly, this current analysis dissects the velocity of a locus into effects mediated by internal contacts that restrict intrinsic movement and external forces that act on the chromatin fiber. We note that in mammalian cells, cytoplasmic microtubules linked to the nucleus through SUN domain proteins (or in yeast the SPB) can increase nuclear oscillations, generating a detectable degree of chromatin motion. Whereas cytoplasmic microtubules alter chromatin movement in mammalian cells during DNA damage, it appears to be untrue in yeast. Rather, the depolymerization of the actin cytoskeleton in yeast allowed us to differentiate such oscillations from the alterations at a specific site, that reduce tethering, enhance anomalous movement, decompact the polymer and lead to extrusion of the damaged site. These latter changes required the nucleosome remodeler INO80, and can be modeled and simulated as reduced monomer contact within a *β*-polymer (Fig. 3–5).

### 4.1 Forces acting on chromatin are reduced after DSB induction

The changes arising from the decondensation of chromatin surrounding the DSB should be independent of nuclear organization and cell type, because they rely only on local changes within the chromatin fiber. On the other hand, general HR search time will depend strongly on nuclear organization, the density of chromatin and the presence or absence of chromosomal territories. Based on both live image analysis [11] and the present computational simulations, we argue that the search time for interaction between a break and a homologous donor sequence depends on the diffusion constant, the length of the polymer, local forces acting on the DSB, the position of the interacting monomer along the polymer chain, and the position of the DSB relative to the nuclear envelope. In the case of subnuclear positioning and INO80-induced modifications of break-proximal nucleosomal organization, experimental data confirm their impact on HS [4, 39].

We have shown here that the drop in local chromatin interactions at the site of the DSB provokes a change in chromatin organization that leads to the extrusion of the break from the chromatin domain. The extrusion of a DSB massively accelerates the search process (Fig. 4). The effect is further amplified when both the DSB and the homologous target shift to the surface of their respective domains (Fig. 5). This extrusion of damages from a chromatin domain could facilitate not only HR, but NHEJ or other repair pathways, by increasing accessibility of the damage to repair factors [40]. This may also occur at uncapped mammalian telomeres, where local movement was shown to facilitate telomere-telomere fusions in a 53BP1-dependent manner [41, 42]. Finally, as discussed below, DSB extrusion appears to happen at damage within heterochromatin, as demonstrated in Drosophila [43], and in the yeast ribosomal DNA [44].

### 4.2 Chromatin decondensation can accelerate the search process

Using simulations of a *β*-polymer model, we accounted for long-range interactions between monomers and show that the release of inter-monomer forces can also be associated with the relaxation/decondensation of chromatin (Fig. 3). Our simulations show that chromatin relaxation accelerates HS, as it also reduces the screening barrier that is imposed when the broken locus remains inside chromatin. A monomer located within a domain must contact a template located inside a compacted polymer and the surrounding monomers generate an effective potential barrier that needs to be overcome for the encounter to occur. This potential barrier is due to a local effect from all interacting monomers [45] (Fig.5). This situation is similar to the classical chemical reaction theory where a catalyst reduces the activation potential barrier, accelerating the reaction.

### 4.3 Extrusion of a DSB from a chromatin territory: a novel step in HR

When a DSB is repaired by HR, the dsDNA is resected and nucleosomes are evicted for up to several kb around the break [22]. This does not occur readily in heterochromatin [46]. Indeed, in heterochromatin in Drosophila and mammals, and in the rDNA locus in yeast, DSBs must relocate away from the heterochromatic domains for repair to occur [43, 44]. The change in motion experienced by a locus after break induction reflects an increase in diffusion coefficient, anomalous exponent and confinement radius, and, based on numerical simulations, damage extrusion from the interior to the periphery of a chromatin domain (Fig. 5). These changes are predicted to arise from a reduction in internal chromatin constraints, and we proposed that nucleosome remodeling contributes to this, since the loss of Arp8, an essential component of the INO80 nucleosome remodeling complex, impedes loss of local tethering forces at a DSB [11]. Histone tail acetylation may also help overcome local tethering to facilitate DSB extrusion. This latter is a novel prediction that would particularly facilitate repair in a condensed chromatin environment. A recent study has looked at chromatin folding changes within different epigenetic states, and the next step would be to characterize these dynamical changes [47].

There are a number of repair scenarios in which extrusion of a locus to the surface of a domain might be important, in addition to the repair of DSB in heterochromatin by HR. In an earlier study, [41] showed that de-protected telomeres become more mobile and that this enhanced mobility correlates with telomere-telomere fusions. Both the fusions and the movement are dependent on the binding of 53BP1, a factor that promotes NHEJ. It is possible, that the extrusion of damage from a compacted chromatin environment, also enables access to other repair machineries such as those mediating telomere-telomere ligation. Nonetheless, in a crowded environment such as the nucleus, self-avoiding interactions have a dominant effect on the MFET during repair by HR. These interactions can be regulated by changing monomer density, monomer-monomer interaction, and dynamics.

We have shown here that loop, forces and cross-linkers used in polymer models can explain many observations related to single particle trajectories of DNA loci before and after DSB induction. This approach is only the first step toward understanding HS through polymer physics. This goal will require a multi-scale approach that integrates both fine and coarse-grain behavior of the chromatin fiber, and Chromosome Conformation Capture data. Precise details about local protein organization, strand invasion and nucleosome eviction must be integrated in a model to uncover novel features of HR and other encounter mechanisms in the nucleus.

## 5 Methods

### 5.1 Randomly cross-linked (RCL) polymer

The RCL polymer [20] is composed of *N* monomers *R* = [*r*_1_, *r*_2_, …, *r*_*N*_] in dimension *d* connected sequentially by harmonic springs, and additional *N*_*c*_ springs connecting randomly chosen monomer pairs in each realization 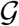. The added connectors mimic the effect of binding molecules (CTCF) and their random position over all realizations reflects the heterogeneity in chromatin structure over cell population. The spring potential of the RCL polymer is composed of the potential of springs of the linear backbone and that of the added connectors, given in the quadratic form

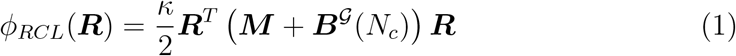

where *κ* is the spring constant, ***M*** is the connectivity matrix for the linear backbone of the polymer (Rouse matrix), given by

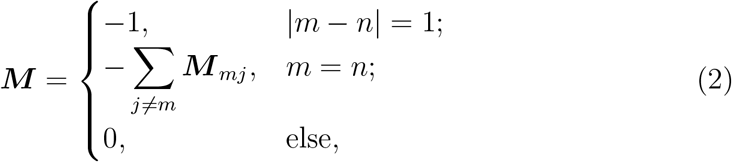

and ***B***(*N*_*c*_) is the added connectivity matrix

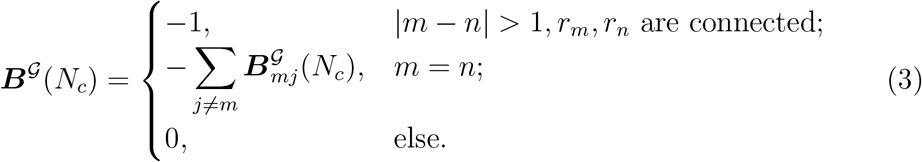

The dynamics of monomers ***R*** of the RCL polymer is given by

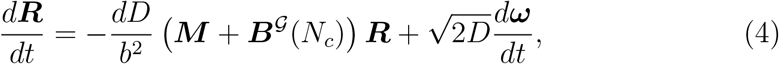

where *D* is the diffusion constant, *b* is the standard deviation of springs’ length, *ω* are *d*-dimensional Brownian motions,

### 5.2 Simulations of a DSB in the RCL polymer

To examine the effect of a single DSB on the anomalous exponent *α*, the apparent diffusion coefficient *D*, the coefficient of constraint *k*_*c*_, and the length of constraint *L*_*c*_, we perform numerical simulations of the RCL polymer 4 of *N* = 100 monomers in dimension *d* = 3, and *N*_*c*_ = 50 connectors between non nearest neighboring monomer pairs. For each realization 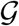, we randomize the position of the *Nc* added connectors and we first simulate system 4 up to its relaxation time *τ*_*R*_ defined by

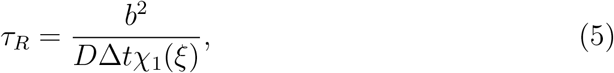

with Δ*t* = 0.0033 *s*, where *χ*_1_ is the first non-vanishing eigenvalue of the connectivity matrix (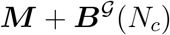) in realization 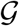, which we compute numerically. In practice, the number of steps to relaxation was in the range of tens of thousands of steps (33s-2minutes). At time *τ*_*R*_, we induce a single DSB between the two middle monomers *r*_50_, *r*_51_ by removing the spring connectors between them. We discard realizations in which the two parts of the chain where completely disconnected following DSB. We then continue simulations for an additional *N*_*s*_ = 10000 steps to a total of 33 seconds post DSB.

We study three scenarios where connectors are discarded following DSB: 1) *N*_*r*_ < *N*_*c*_ added connectors are removed uniformly at random, and 2) *N*_*r*_ nearest connectors (in spatial distance from the average position of monomers *r*_50_ and *r*_51_ at time *τ*_*R*_) are removed around the DSB, and 3) all cross-links at a distance *d*. We perform 1000 simulations for each *N*_*r*_ ∈ [0, 45] number of connectors removed. Parameters used in simulation are summarized in Table 1.

### 5.3 Simulations of loci trajectories following DSB

We now describe how we evaluated from RLC-polymer simulations the four parameters of interest (anomalous exponent *α*, Length of confinement *Lc*, apparent diffusion coefficient *D*, effective spring coefficient *k*_*c*_) from trajectories of monomers *m* = 1, … *N*.

#### 5.3.1 Computing the anomalous exponent α

To estimate the mean-square-displacement of each monomer *m* = 1, …, 100 located at position *r*_*m*_, we use the formula

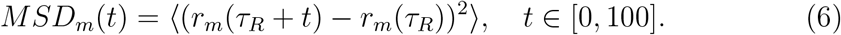

We then fitted the function 6 with

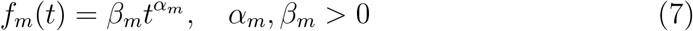

to extract the value of the anomalous exponent *α*_*m*_ of monomer *m*. We waited an equilibrium *τ*_*R*_ in the range of 20s to few minutes.

#### 5.3.2 Computing the Length of confinement *Lc*

The length of confinement *L*_*c*_ of monomer *r*_*m*_ in each realization is estimated by averaging over *N*_*s*_ = 3000 simulation steps. We use the empirical estimator [48]

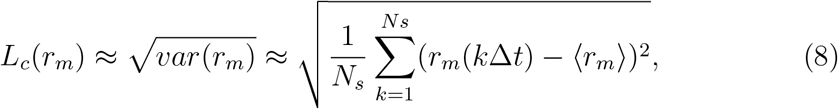

where 〈*r*_*m*_〉 is the average position of monomer *m* throughout the simulation.

#### 5.3.3 Computing the apparent diffusion coefficient *D*

The apparent diffusion coefficient of monomer *r*_*m*_ is estimated by the formula

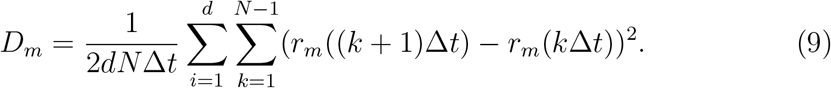

The parameter are defined in table 1.

#### 5.3.4 Computing the effective spring coefficient *k*_*c*_

The force acting on each monomer due to confinement is modeled as a harmonic spring force [49]. The associated spring constant is estimated by the empirical formula

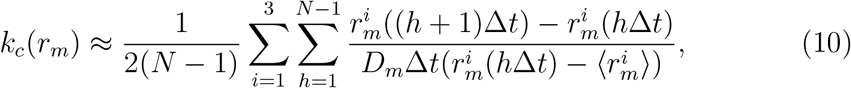

where the superscript indicates the dimension of the space. In practice [48], the quantity 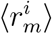 is computed by averaging over trajectories. The diffusion coefficient *D*_*m*_ can be computed by using formula 9. In practice, to avoid dividing by small number when the monomer position fluctuates around the mean position, we use a linear regression to estimate the slope for 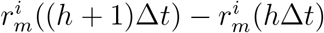 versus 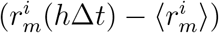.

##### Modeling and simulations with Rouse polymer

A Rouse polymer is a collection of monomers positioned at ***R***_*i,n*_ (*n* = 1, 2, …*N, i* = *a, b*), moving with a random Brownian motion coupled to a spring force originating from the nearest neighbors. The potential energy is

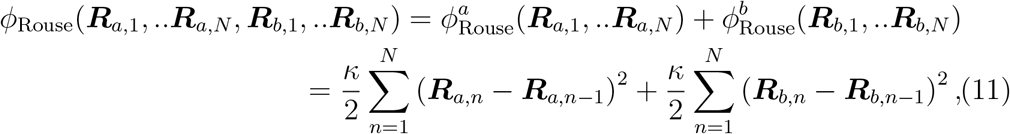

where the spring constant *κ* = 3*k*_*B*_*T/b*^2^ is related to the standard-deviation *b* of the distance between adjacent monomers [50] with *k*_*B*_ the Boltzmann coefficient and *T* the temperature. In units of *k*_*B*_*T*, we have *κ* = 3/*b*^2^ and *D* = 1/*γ*, where *γ* is the friction coefficient. In the Smoluchowski’s limit of the Langevin equation [31], the dynamics of monomer ***R***_*i,n*_ is

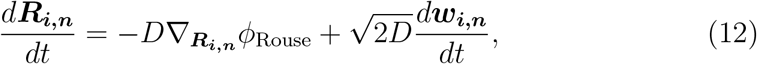

for *n* = 1, …, *N* and *i* = *a, b*, and ***w***_*i,n*_ are independent three-dimensional white noise with mean zero and variance 1. We often focus on two monomers located on two different Rouse polymers *a* and *b* with the same length (*N*_*a*_ = *N*_*b*_ = *N*). The search time 〈*τ*_*e*_〉 between two monomers is the mean time for the two monomer *n*_*a*_, *n*_*b*_ to come into a distance *ε* < *b*

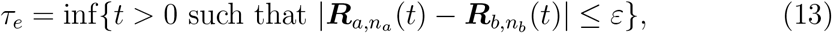

while polymers are restricted in a confined domain.

##### Implementing excluded volume interactions

We use other polymer models by adding interactions such as bending elasticity, which accounts for the persistence length of the polymer and Lennard-Jones forces (LJ), describing self-avoidance of each monomer pairs. To account for thee LJ forces, we use the potential energy defined by

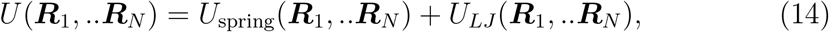

where the spring potential is

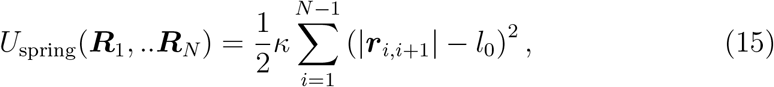

where *l*_0_ is the equilibrium length of a bond, 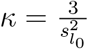 and *s*_*l*0_ is the standard deviation of the bond length. We chose the empirical relation *s*_*l*0_ = 0.2*l*_0_. The Lennard-Jones potential is

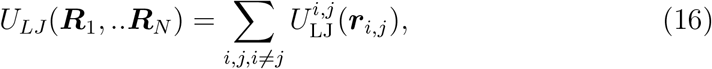

with ***r***_*i,j*_ = ***R***_*i*_ − ***R***_*j*_ and

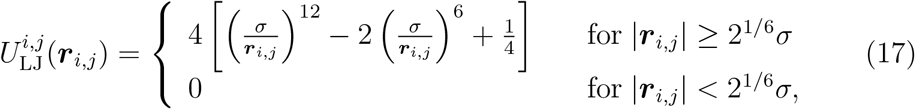

where *σ* is the size of the monomer. With the choice *l*_0_ = 2*σ*, the springs which materialize bonds, cannot cross each other in stochastic simulations using eq. 12 with potential 14. We do not account here for bending elasticity. Finally, we used Euler’s scheme to generate Brownian simulations. At impenetrable boundary, each rigid monomer is reflected in the normal direction of the tangent plane.

##### The *β*−polymer model

To account for the chromatin dynamics, we use the generalized polymer model called the *β*-model [19], which accounts for anomalous diffusion behavior with an exponent in the range of]0 − 0.5], as measured for yeast in vivo chromatin locus [51]. We use *β*-polymer model where all monomers are connected through a quadratic potential defined by

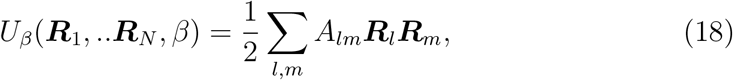

with coefficients

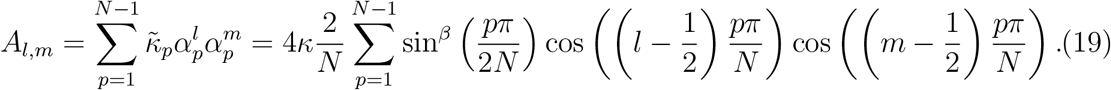

and

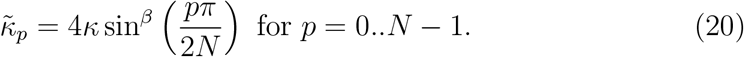

In such model, the strength of interaction *A*_*l,m*_ decays with the distance |*l − m|* along the chain. By definition, 1 < *β* < 2 [19] and the Rouse polymer is recovered for *β* = 2, for which only nearest neighbors are connected.

##### Steady-state configuration of a *β* polymer with Lennard-Jones forces

To account for changes in the chromatin structure and to avoid inter-penetration of the chromosome, we added the classical Lennard-Jones (LJ) interactions to the polymer model (see Experimental Procedures). We considered a polymer of length *N* = 33 with a coefficient *β* = 1.5, where all monomers are highly connected. We point out here that with the addition LJ-interaction forces, the anomalous exponent *α* does not satisfies the previous relation *α* = 1 − 1/*β*. Thus, we estimated numerically the anomalous exponent of a self-avoiding polymer with *β* = 1.5 and found *α* = 0.52 for a tagged monomer (data not shown), a prevalent experimental value.

